# Improving the Safety of Cultured Meat Using Basal Medium Prepared using Food Ingredients

**DOI:** 10.1101/2022.03.28.486069

**Authors:** Takanori Kanayama, Keita Tanaka, Haruna Shimada, Koji Ichiyama, Sadaharu Higuchi, Ikko Kawashima

## Abstract

Currently, meat production involves environmental issues and high costs; one solution to address this may be to produce meat via cell culture. Therefore, there is an urgent demand for the development of cell culture methods that replace raw materials with food materials to produce cultured meat as food. In this study, we designed food-grade-Dulbecco’s Modified Eagle’s Medium (FG-DMEM); when the cells cultured in FG-DMEM and regular DMEM were compared, there was no significant difference in cell growth or differentiation. Therefore, FG-DMEM can be used for research and safe production of cultured meat.

## Main text

The demand for meat is increasing with the growth in population. Most meat products are produced by livestock farming and fishing (wild fishing/fish farming). However, the environmental costs of this production are high; thus, alternative methods to achieve sustainable meat development are urgently needed.

Cultured meat has attracted attention as an alternative that can reduce the environmental costs of livestock production^1^. Meat production via cell culture can reduce energy consumption by 7–45%, greenhouse gas emissions by 78–96%, land use by 99%, and water consumption by 82–96%^2^. Recently, the Singapore Food Agency (SFA), has given approval for commercial sale of cell-based meat products for local consumption by US firm Eat Just. The SFA said it was the first country to pioneer a regulatory framework for novel foods and approve the commercial sale of cell-based meat^3^. Currently, the production of cultured meat is increasing worldwide. Cultured meat is obtained by culturing animal cells and differentiating them by adding nutrients and growth factors to the medium^4^. However, in vitro, meat is cultured in a basal medium prepared using chemical reagents used for cell culture. To scientifically verify and simulate the production of cultured meat as a food product, a basal medium consisting only of food/food additives must be developed. Therefore, a food-grade medium was designed based on Dulbecco’s Modified Eagle’s Medium (DMEM), with all components of the DMEM replaced with food/food additives. Food-grade (FG)-DMEM is composed of ingredients that have been consumed as food, ensuring the safety and enabling regulatory clearance (such as for Generally Recognized as Safe rules) in the United States.

In this study, mouse skeletal muscle-derived cells (C2C12) and bovine skeletal muscle-derived primary cells (BSMCs) were used because the edible portion of meat is the muscle. The cells were cultured in FG-DMEM and DMEM to confirm whether the developed medium is suitable for cell culture. The FG-DMEM composition was designed to be the same concentration as DMEM after dissolution (Table 1 and Supplementary Table 1). Also, the amount added was calculated according to the lower value of the purity range (Supplementary Table 2). As the purity of food ingredients has not been clearly defined, ingredient levels were set according to the minimum contents indicated for food ingredients (for detailed methods, see the supplementary information). Additionally, we substituted materials for which food-grade alternatives were not available, including Fe(NO_3_)_3_·9H2O for FeCl_3_·6H_2_O, L-arginine HCl for L-arginine, L-histidine HCl·H_2_O for L-histidine, and choline chloride for glycerophosphatidylcholine.

**Table 1.** Composition table of culture medium using food ingredients. This composition table describes the types of raw materials and their contents in 1 L of culture medium. These food materials are at least listed on the positive list as food ingredients in Japan.

| Components | (mg/L) | Components | (mg/L) |
| --- | --- | --- | --- |
| ***Inorganic components*** |  | ***Amino acids*** |  |
| CaCl<sub>2</sub> · 0-2H<sub>2</sub>O | 378.44 | L-Arginine · HCl | 70.88 |
| KCl | 404.04 | L-Cystine | 49.33 |
| MgSO<sub>4</sub> · 7H<sub>2</sub>O | 202.01 | L-Glutamine | 595.88 |
| NaCl | 5509.41 | Glycine | 30.46 |
| NaHCO<sub>3</sub> | 3737.37 | L-Histidine | 31.72 |
| NaH<sub>2</sub>PO<sub>4</sub> · 2H<sub>2</sub>O | 144.23 | L-Isoleucine | 107.14 |
| ***Trace element*** |  | L-Leucine | 107.14 |
| FeCl<sub>3</sub> · 6H<sub>2</sub>O | 0.07 | L-Lysine · HCl · 2H<sub>2</sub>O | 148.98 |
| ***Vitamins*** |  | L-Methionine | 30.46 |
| Ca Pantothenate | 4.13 | L-Phenylalanine | 67.01 |
| α-GPC | 7.51 | L-Serine | 42.86 |
| Folic acid | 4.08 | L-Threonine | 96.94 |
| i-Inositol | 7.42 | L-Tryptophan | 16.33 |
| Niacinamide | 4.04 | L-Tyrosine | 73.05 |
| Pyridoxine · HCl | 4.08 | L-Valine | 95.92 |
| Riboflavin | 0.41 | ***Other components*** |  |
| Thiamine · HCl | 4.08 | D-Glucose monohydrate | 4999.88 |

FG-DMEM was dissolved in ultrapure water, and the pH was adjusted using food-grade NaOH and food-grade HCl. The pH of FG-DMEM was 7.81 ± 0.04 (N = 3), which was close to that of control DMEM (pH 7.78 ± 0.02). The osmotic pressure was the same for both FG-DMEM (332.67 ± 5.51 mOsm/KgH_2_O) and DMEM (330.67 ± 2.52 mOsm/KgH_2_O).

First, the effect of FG-DMEM on cell proliferation was examined. C2C12 and BSMCs grown in FG-DMEM and DMEM were compared at different passage numbers. The C2C12 cells were passaged twice for acclimation in FG-DMEM and DMEM. No difference was observed for each medium until at least the fifth passage (Fig. 1A). Similarly, there was no significant difference in BSMCs grown in FG-DMEM and DMEM until at least the fifth passage (Fig. 1B). These results suggest that FG-DMEM can be used as an alternative to DMEM for inducing cell proliferation.

**Fig. 1.**
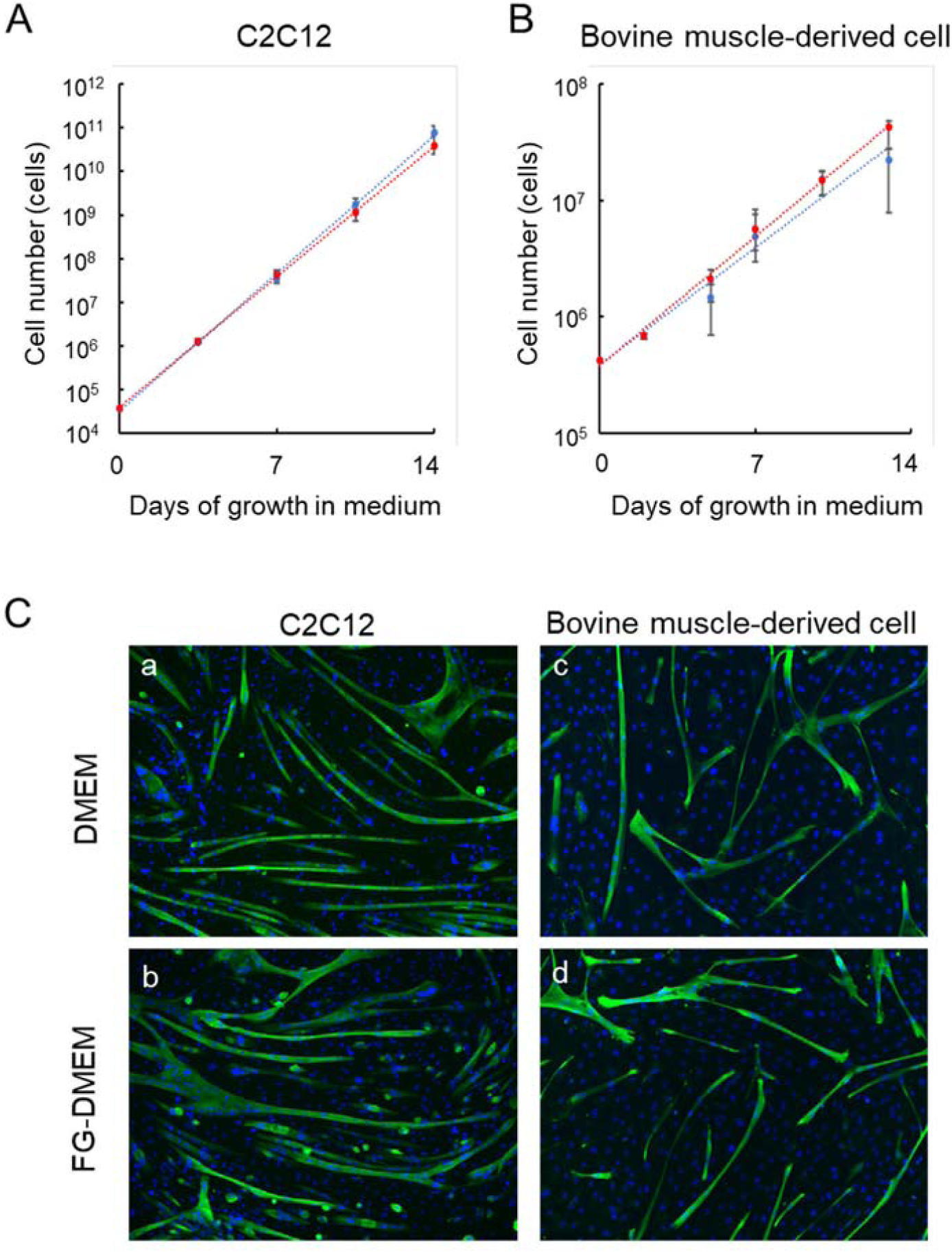
Effect of FG-DMEM on the proliferation and differentiation of skeletal muscle-derived cells. C2C12 (A) and bovine muscle-derived cells (B) were cultured up to five passages in DMEM and FG-DMEM with 10% FBS, to count the number of cells. The culture was passaged every three days, and the number of cells was counted each time with an automated cell counter. C2C12 and bovine muscle-derived cells were cultured in DMEM and FG-DMEM with house serum and observed for myosin heavy chain staining. C2C12(B-a,b) were observed after seven days of differentiation induction culture, and bovine muscle-derived cells(B-c,d) were observed after three days of differentiation induction culture.

Next, the effects of FG-DMEM on inducing the differentiation of cultured C2C12 and BSMCs were examined. C2C12 and BSMCs were grown in FG-DMEM or DMEM for five passages. C2C12 fused and formed myotubes in FG-DMEM and DMEM at day 7 (Fig. 1C). The myotube fusion index for cells cultured in FG-DMEM was not significantly different (5.71 ± 0.75%) from that of cells grown in DMEM (4.11 ± 0.36%). Next, we induced the differentiation of BSMCs in each medium and found that myotubes formed at day 3 in each group. Furthermore, the fusion index for cells cultured in FG-DMEM was not significantly different (7.25 ± 1.88%) from that of cells cultured in DMEM (8.52 ± 3.02%). These results indicate that a culture medium prepared using food materials can induce myotube formation as effectively as DMEM.

Our results indicate that FG-DMEM is promising for use in cultured meat production. The culture medium developed in this study can also be used for scientific validation and research of cultured meats. Culture of cells in FG-DMEM showed results similar to those of culture in DMEM. Although FG-DMEM is as useful as DMEM basal medium, cultures in FG-DMEM can have different cell phenotypes compared to DMEM cultures. In conclusion, this study provides a foundation for using food-derived raw materials for meat production via cell culture and for improving the safety of cultured meat. However, in the cell acclimatisation experiments (data not shown), subtle differences in the amount and composition of ingredients in each medium can affect cultured cell phenotype. Further verification of our results is necessary.

In addition to DMEM, various other basal media are used for skeletal muscle culture. Studies are needed to determine whether these media can be used in combination with food ingredients.

## Methods

### Preparation of food-grade basal medium

The food-grade components of DMEM are listed in Supplementary Table 1 and Supplementary Table 2. These reagents were dissolved in ultrapure water (PURELAB Chorus 1 Complete; Bio Medical Sciences, Tokyo, Japan) as shown in Table 1. Then, the pH was adjusted with NaOH and the solution was sterilised with a 0.22-μm filter (TPP Techno Plastic Products, Trasadingen, Switzerland) and refrigerated until use.

### C2C12 cell culture

The C2C12 mouse myoblast cell line was maintained in DMEM (FUJIFILM Wako Pure Chemical, Osaka, Japan) containing 1% penicillin-streptomycin-amphotericin B (FUJIFILM Wako Pure Chemical) and 10% (v/v) foetal bovine serum (FBS; FUJIFILM Wako Pure Chemical) at 37°C and 5% CO_2_. Differentiation was induced by changing the medium supplement from 10% FBS to 2% horse serum (Thermo Fisher Scientific, Waltham, MA, USA).

### Preparation and cell culture of BSMCs

Fresh post-mortem muscle sections were obtained from bovine cheeks (Tokyo Shibaurazoki, Tokyo, Japan) and treated with chlorhexidine gluconate solution (Sumitomo Dainippon Pharm, Japan) and mechanically dissociated using collagenase type IV to obtain BSMCs, which were cultured in the same manner as C2C12 cells.

### Proliferation assay

The cells were seeded in 60-mm dishes with DMEM or FG-DMEM, containing 10% FBS for C2C12 cells and 20% FBS for BSMCs. The seeding densities were 1.2 × 105 C2C12 cells and 2.2 × 105 bovine muscle-derived cells. After harvest, the cells were trypsinised and counted with a haemocytometer (AR BROWN, Tokyo, Japan) each day.

### Immunofluorescence staining

After induction of myotube differentiation, the cells were stained using standard methods^5^. A mouse anti-myosin heavy chain antibody was used as primary antibody (1:200; Sigma Aldrich, St. Louis, MO, USA) and Alexa 488 donkey anti-mouse IgG antibody as secondary antibody (Life Technologies, Carlsbad, CA, USA).

### Image analysis

Images were processed using ImageJ software (NIH, Bethesda, MD, USA). All measurements were performed manually. The fusion index was calculated dividing the number of nuclei in Desmin- or MyHC-positive cells by the number of nuclei in the image. For every sample, at least five images were evaluated.

### Statistical analysis

Student’s t-test was performed using Excel software (Microsoft, Seattle, WA, USA). Two-way analysis of variance was performed in R. Data are presented as the means ± standard error of the mean.

## Acknowledgements

We thank Yuki Hanyu, Keita Fukumoto, Keisuke Igarashi and Minami Sone (IntegriCulture, Inc.) for providing useful comments and editing the manuscript. This work was supported by a Grant-in-Aid for JST MIRAI (No. 18077323) from the Japan Society for the Promotion of Science (to I.K.).

## Autor contributions

I.K. designed the research and wrote the manuscript. T. K and K.T designed and performed the experiments. H.S and K.I performed the experiments. S.H and I.K contributed to the review and editing of the manuscript. All authors critically revised and commented on the report and approved the final manuscript.

---

**Supplementary Table 1.**
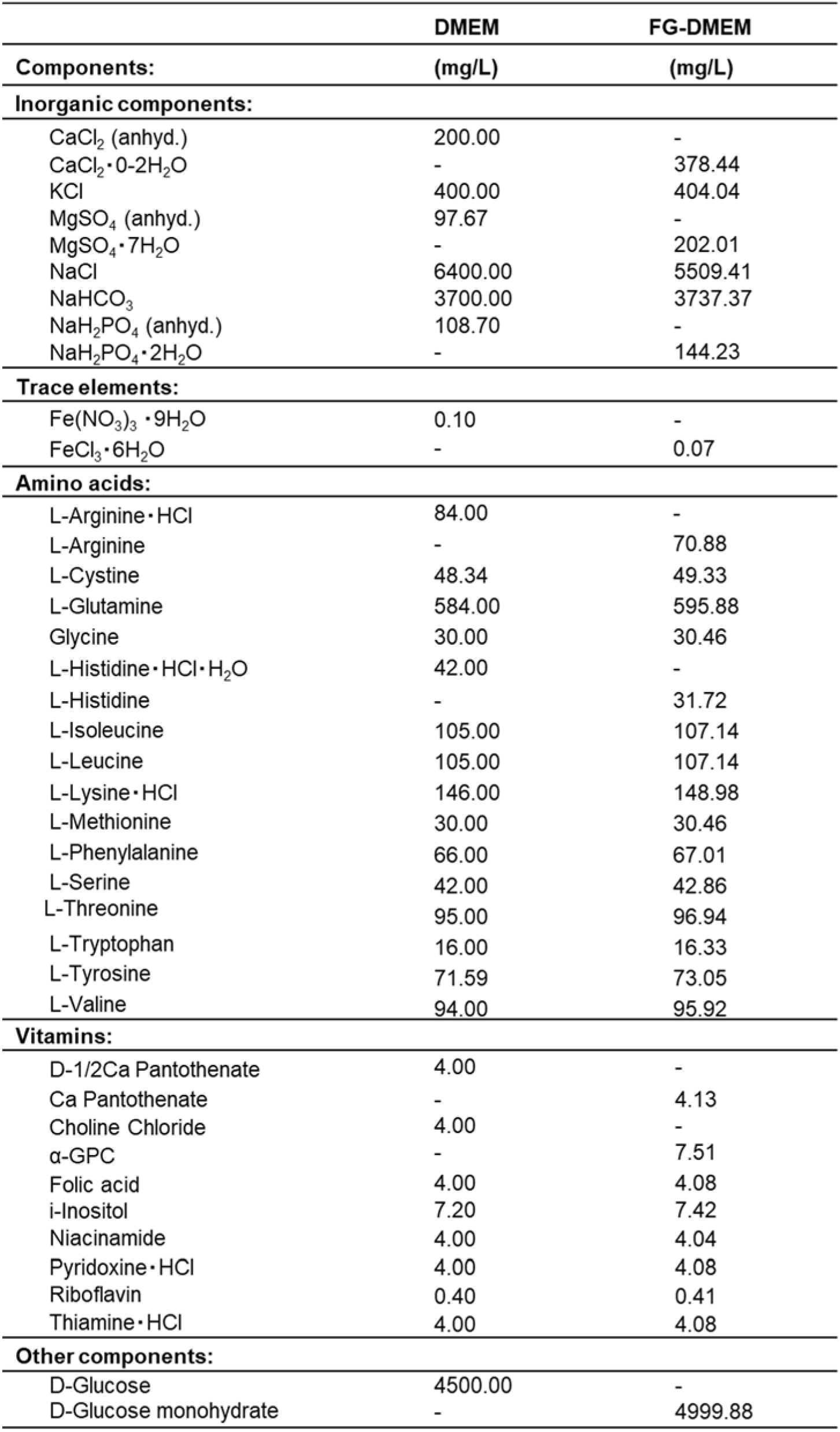
Comparison of ingredients and quantities of FG-DMEM and DMEM.

|  | DMEM | FG-DMEM |
| --- | --- | --- |
| Components: | (mg/L) | (mg/L) |
| **Inorganic components:** |  |  |
| CaCl<sub>2</sub> (anhyd.) | 200.00 | - |
| CaCl<sub>2</sub> • 0-2H<sub>2</sub>O | - | 378.44 |
| KCl | 400.00 | 404.04 |
| MgSO<sub>4</sub> (anhyd.) | 97.67 | - |
| MgSO<sub>4</sub> • 7H<sub>2</sub>O | - | 202.01 |
| NaCl | 6400.00 | 5509.41 |
| NaHCO<sub>3</sub> | 3700.00 | 3737.37 |
| NaH<sub>2</sub>PO<sub>4</sub> (anhyd.) | 108.70 | - |
| NaH<sub>2</sub>PO<sub>4</sub> • 2H<sub>2</sub>O | - | 144.23 |
| **Trace elements:** |  |  |
| Fe(NO<sub>3</sub>)<sub>3</sub> • 9H<sub>2</sub>O | 0.10 | - |
| FeCl<sub>3</sub> • 6H<sub>2</sub>O | - | 0.07 |
| **Amino acids:** |  |  |
| L-Arginine • HCl | 84.00 | - |
| L-Arginine | - | 70.88 |
| L-Cystine | 48.34 | 49.33 |
| L-Glutamine | 584.00 | 595.88 |
| Glycine | 30.00 | 30.46 |
| L-Histidine • HCl • H<sub>2</sub>O | 42.00 | - |
| L-Histidine | - | 31.72 |
| L-Isoleucine | 105.00 | 107.14 |
| L-Leucine | 105.00 | 107.14 |
| L-Lysine • HCl | 146.00 | 148.98 |
| L-Methionine | 30.00 | 30.46 |
| L-Phenylalanine | 66.00 | 67.01 |
| L-Serine | 42.00 | 42.86 |
| L-Threonine | 95.00 | 96.94 |
| L-Tryptophan | 16.00 | 16.33 |
| L-Tyrosine | 71.59 | 73.05 |
| L-Valine | 94.00 | 95.92 |
| **Vitamins:** |  |  |
| D-1/2Ca Pantothenate | 4.00 | - |
| Ca Pantothenate | - | 4.13 |
| Choline Chloride | 4.00 | - |
| α-GPC | - | 7.51 |
| Folic acid | 4.00 | 4.08 |
| i-Inositol | 7.20 | 7.42 |
| Niacinamide | 4.00 | 4.04 |
| Pyridoxine • HCl | 4.00 | 4.08 |
| Riboflavin | 0.40 | 0.41 |
| Thiamine • HCl | 4.00 | 4.08 |
| **Other components:** |  |  |
| D-Glucose | 4500.00 | - |
| D-Glucose monohydrate | - | 4999.88 |

**Supplementary Table 2.**
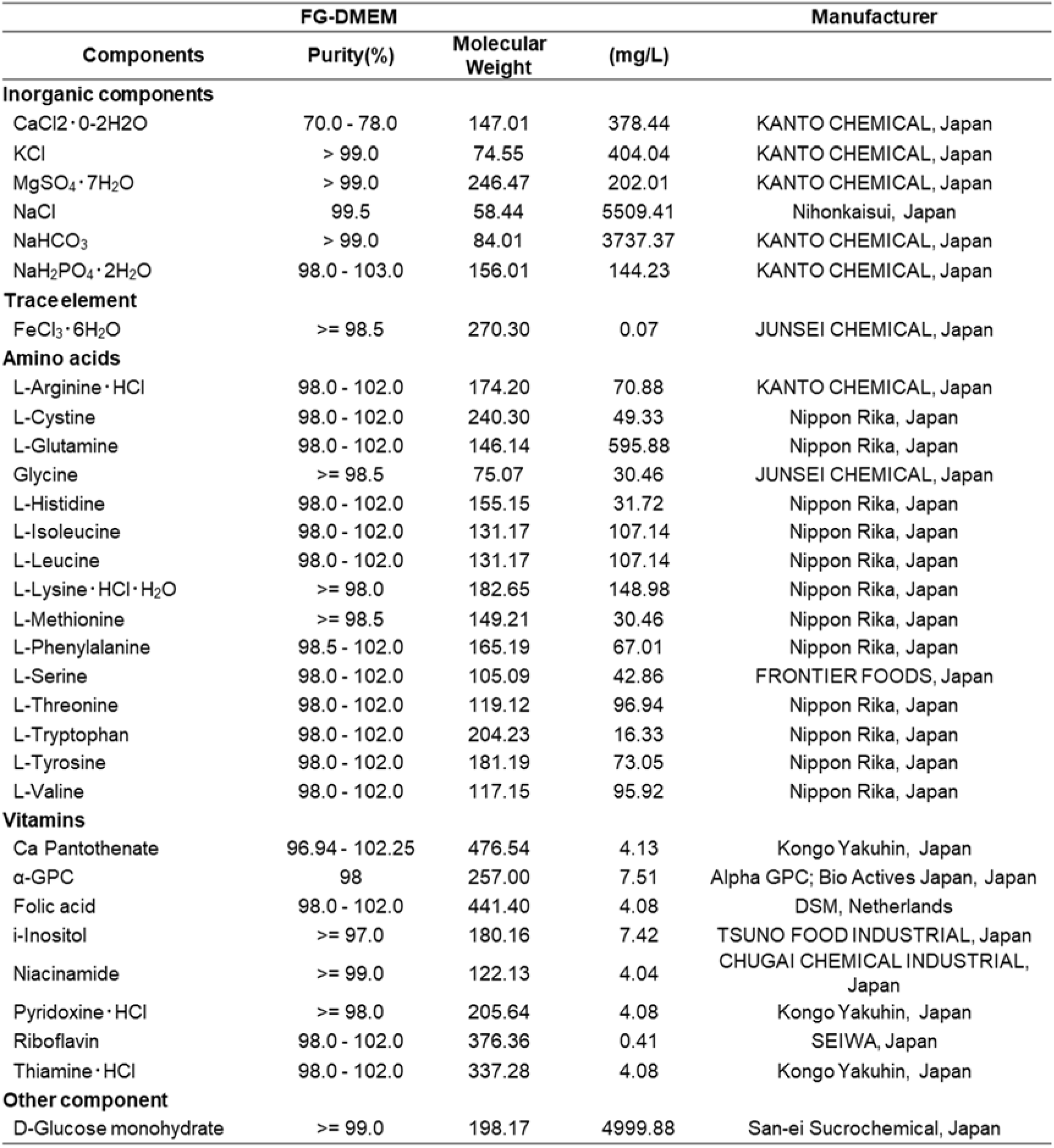
The table describing food ingredients, purity and quantity of raw materials. To add the same amount of raw material as the composition of DMEM, the amount added was calculated according to the lower value of the purity range. The values in the table are the amounts of raw materials needed to make 1 L of FG-DMEM.

